# Dimerization Mechanism of HIV-1 RNA Hairpins to Extended Duplex Structures

**DOI:** 10.1101/2025.10.22.683870

**Authors:** Dibyendu Mondal, Sk Habibullah, Govardhan Reddy

## Abstract

Genomic RNA (gRNA) dimerization is essential for retroviral replication. In the gRNA of human immunodeficiency virus (HIV-1), the hairpin-like dimerization initiation sequence (DIS) forms a kissing-complex (KC) with the DIS sequence in another gRNA, which later converts into a stable extended-duplex (ED). Using coarse-grained simulations, we mapped the transition of HIV-1 DIS RNA hairpins (HPs) to ED and identified multiple intermediates beyond the KC. The KC has an anionic pocket stabilized through the condensation of Mg^2+^ ions. Hence, when only K^+^ ions are present at physiological levels, the HPs to ED transitions occur through a different dominant pathway devoid of the KC. We also observed purine base flipping near KC hydrogen bonds, revealing the population of a sub-ensemble of intermediates. The proposed dimerization mechanism of HPs to ED, along with the sub-ensemble of KC conformations and its anionic pocket, provides a strategic framework for designing specific retroviral drugs targeting this pathway.

**For Table of Contents Use Only:** 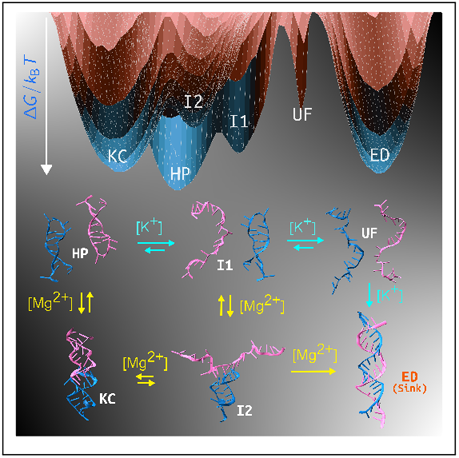

---

RNA dimerization is critical to the life cycle of retroviruses as it facilitates the packaging of two genomic RNA copies into a virion.^1,2^ In retroviruses like human immunodeficiency virus (HIV) and murine leukemia virus (MuLV), the genomic RNA (gRNA) dimerization is essential for selecting and packaging the correct RNA into new viral particles.^1,3,4^ This process ensures efficient replication, influences RNA translation, supports strand transfer during reverse transcription to maintain genome integrity, and promotes genetic recombination. These functions enhance viral genetic diversity, contributing to viral evolution and adaptability. ^1,5–8^

Due to the importance of RNA dimerization in the survival of retroviruses, extensive experimental and theoretical studies have been conducted to understand its mechanism, using HIV-1 RNA as a model system.^1,9–11^ HIV harbors two homologous copies of its gRNA, which engage in extensive intermolecular interactions, such as the dimer linkage structure (DLS) located within the 5’ untranslated region. ^9,12^ DLS contains a highly conserved sequence known as the dimerization initiation sequence (DIS), which has a short palindromic sequence essential for dimerization.^9,13,14^ Central to this process is the formation of a specific RNA structural motif at DIS, called kissing complex (KC), which initiates the dimer formation. ^9,11,15^

In vitro studies^1,9,10,15–19^ of small fragments of DLS showed that initially it forms a KC, which reorganizes to a stable extended duplex (ED) (Fig. 1a). X-ray crystallographic structures reported the structure of the ED ^19,20^ and polymorphic nature of the KC conformations^15,21^ (Fig. 1b,c). Single-molecule FRET (smFRET) experiments using two 23-nucleotide HIV-1 DIS hairpin constructs (DIS-1 and DIS-2) ^9^ revealed that they partially fuse to form the KC, which transforms into ED via a bent-kissing-complex (bent-KC) intermediate under the strong influence of Mg^2+^ ions (Fig. 1a). NMR studies complemented the KC crystal structures; however, they differ in the specific orientation of the set of bulged purines (G8/A8 and A9) present in the crystal structures. ^10,22–24^ Comprehensive atomistic simulation studies on the KC showed it mostly favors the purines (G8/A8 and A9) in bulged-in conformation, but also predicted that the bulged-out conformations are probable. ^11,25,26^ Similar base flipping has been extensively studied for modules of HIV-1 gRNA other than DIS and reported to be critical for their cellular functioning. ^10,15,18,23,24,27–31^

**Figure 1.**
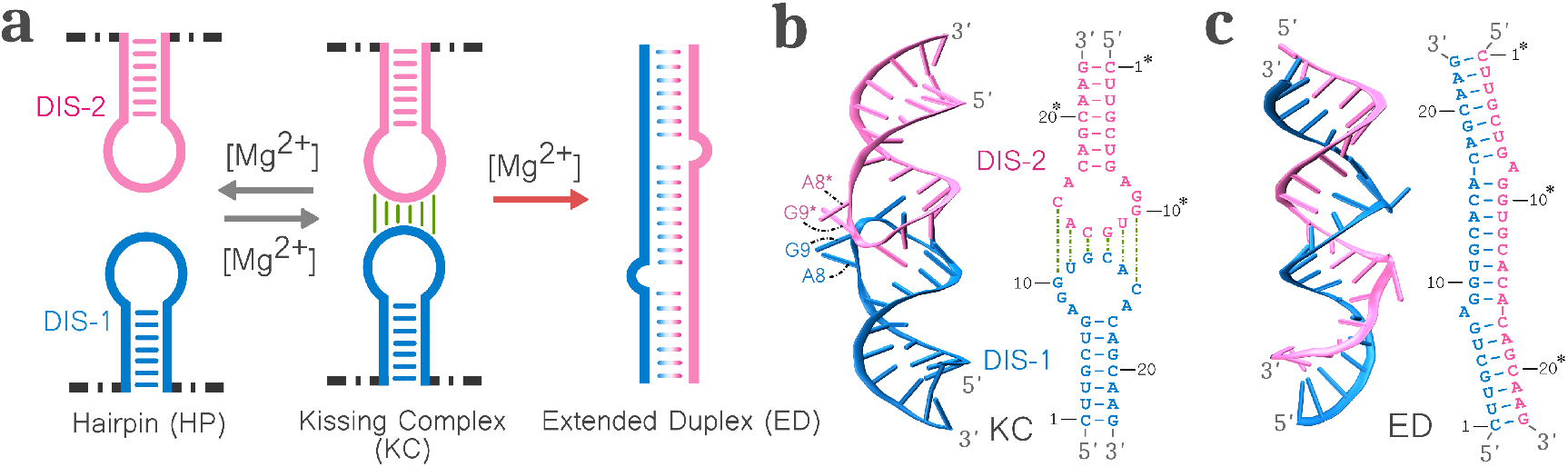
**a.** Dimerization of two DIS hairpin (HP) to extended duplex (ED) via the kissing complex (KC) intermediate. **b**. (left) Crystal structure and (right) secondary structure of KC (PDB ID: 1XPF^15^). The nucleotide indices without and with “*” are for DIS-1 and DIS-2, respectively. The bulged-out residues (A8∗, G9, A8^∗^, and G9^∗^) are marked. **c**. (left) Crystal structure and (right) secondary structure of ED (PDB ID: 462D ^20^).

Despite these extensive studies, the complete thermodynamic landscape illustrating all the critical intermediate states populated during the DIS hairpin (HP) to ED transition is unclear. We further need to understand the mechanism of the reorganization of the hydrogen bond network during the critical KC to ED transition identified in the smFRET experiments. ^9^ We also need to reconcile the subtle structural differences in the orientation of the bulged purines in the KC, as determined by X-ray crystallography and NMR studies. In this work, we present the thermodynamic landscape for the dimerization of a 23- nucleotide DLS construct from HIV-1 subtype A^15,20^ from HP to ED using coarse-grained computer simulations with the three interaction site (TIS) RNA model. ^32–34^ We demonstrate that the DIS HPs populate multiple intermediate states, forming a network that irreversibly leads to the formation of ED. The network connecting the multiple intermediate states is strongly dependent on the concentration of K^+^ ([K^+^]) and Mg^2+^ ([Mg^2+^]). In the presence of Mg^2+^, the dominant pathway for the ED formation is through the KC, and it naturally populates a sub-ensemble of bent conformations. An anionic cavity is present at the core of KC, and the condensation of Mg^2+^ ions in the cavity stabilizes these conformations. Whereas, in the presence of only K^+^, the population of KC is highly diminished and the ED forms through a different dominant route. We further identified the intermediates through which the KC forms the ED by reorganizing the hydrogen bond network. The anionic pocket in the KC was targeted for the development of retroviral drugs.^35^ However, the aminoglycoside molecules identified to bind to the anionic cavity lack specificity, poor penetration in eukaryotic cells, and are toxic at concentrations that exhibit antiviral effects. The multiple sub-ensembles of KC conformations identified in this work could be exploited to enhance the specificity and potency of these molecules.

## Multiple Intermediates are Populated During the Hairpin to Extended Duplex Transition

The dimerization time scale of the DIS HPs to form an ED (PDB ID: 462D^20^) is of the order of seconds.^9^ To overcome this timescale problem, we performed simulations using the TIS RNA model^32,33^ at an elevated temperature (*T* = 333 K) for better conformational sampling ^36,37^ (see Methods). We calculated the fraction of native contacts in the 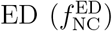 to identify the various intermediates populated during the ED formation (Fig. S1). The state with high 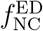 (≈ 0.65) is the ED state. However, we observed that multiple distinct structures are masked in the intermediate states 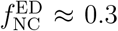 and ≈ 0.02) (Fig. S1) and 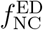 cannot resolve all these distinct states. We find that all transitions to ED from any other state are irreversible, in agreement with previous experiments. ^9^ Therefore, the state ED acts as a sink.

Experiments showed evidence for the population of a kissing-complex (KC) intermediate during the HP to ED transitions,^15,21^ and the crystal structure of the KC is available (Fig. 1b, PDB ID: 1XPF^15^). The KC and ED have 20 and 22 hydrogen-bonded base pairs, respectively. Hence, the KC is stable as an intermediate state. To resolve all the distinct states populated during the HP to ED transition, we also used the fraction of native contacts of the KC, 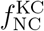. The 2D free energy surface (FES) projected on 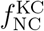 and 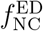 identified all six distinct states populated during the HP to ED transition (Fig. 2a-c, S2, and S3). The conformations of the chains in the five states are: (1) both the chains are unfolded and extended in the basin UF at 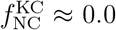 and 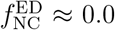(2) both the chains are in the HP conformations in the basin HP at 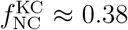 and 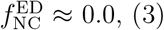 the ED basin is 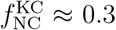 and 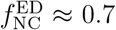 (4) the KC basin is at 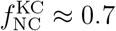 and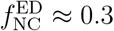, (5) in the basin I1 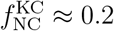 and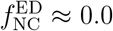, one chain is in the HP conformation and the other chain is in the unfolded extended conformation without any inter-chain interactions, and (6) in the basin I2 at 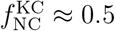 and 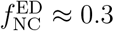 one chain in HP conformation forms inter-chain kissing-loop hydrogen bonding with the other unfolded chain, forming a T-shaped semi-folded structure (Fig. 2d). The KC structure is in equilibrium with structures from other basins before irreversibly forming the ED (Fig. 2a-d, and S1).

**Figure 2.**
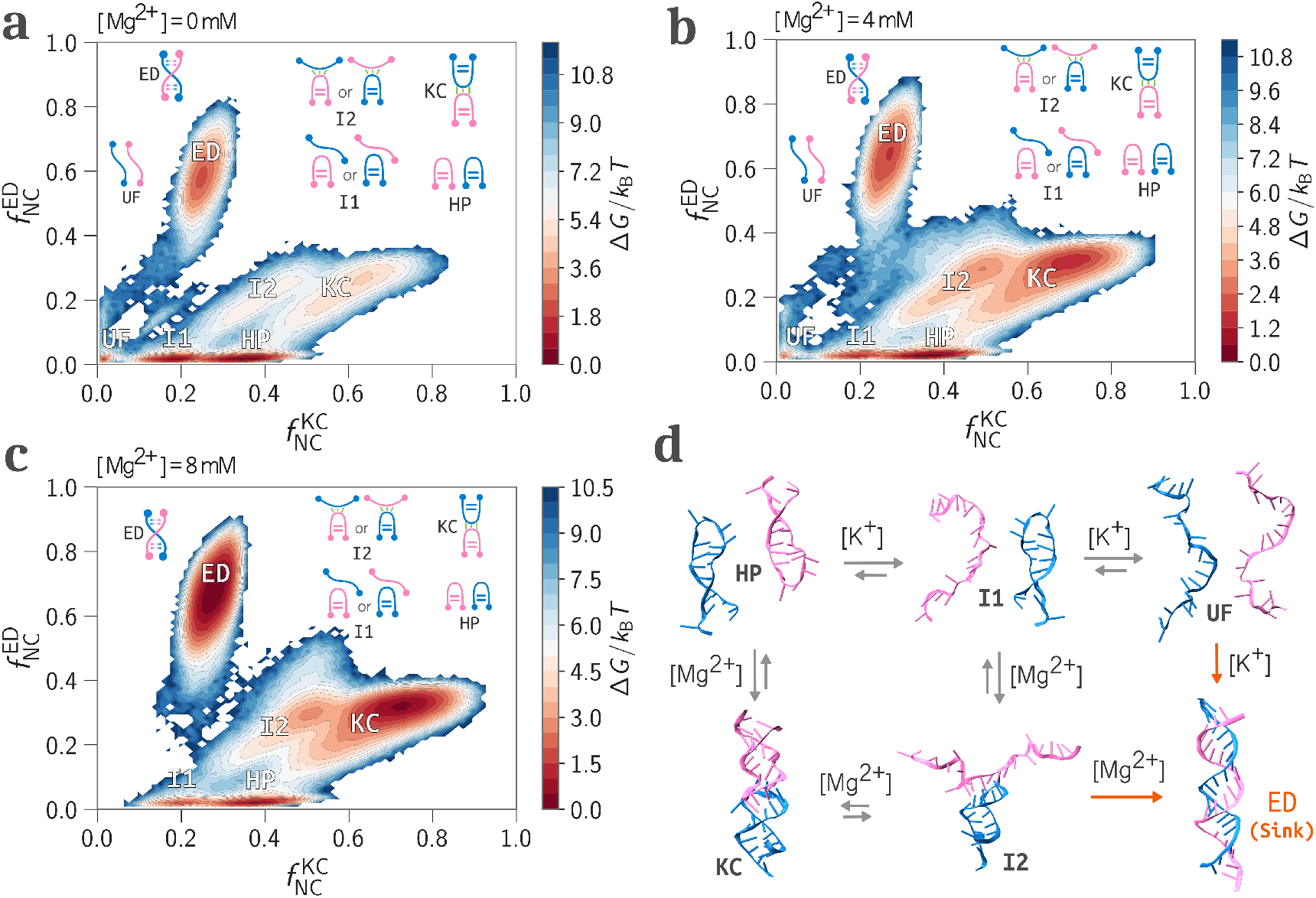
The 2D FES projected onto 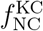 and 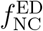 for **a.** [Mg^2+^] = 0 mM **b**. [Mg^2+^] = 4 mM, and **c**. [Mg^2+^] = 8 mM. All have a fixed background [K^+^] = 150 mM. The state ED is a sink. The schematic structures are shown to highlight the structural features in each state. **d**. The network connecting states for the transition from HP to ED. The relative length of the equilibrium arrows highlights whether the particular equilibrium is right or left-shifted for specific ions. The irreversible ED formation is highlighted with the one-way arrow.

## Dimerization Pathways and the Intermediate Population Depend on K^+^ and Mg^2+^ Concentration

The population of intermediate states is strongly influenced by the [K^+^] and [Mg^2+^]. We computed the equilibrium population of different states for [K^+^] = 50, 100, and 150 mM, in the absence of Mg^2+^ (Fig. S4). At low [K^+^] (= 50 mM), the UF state dominates, followed by the I1 state. The population of KC and I2 states is negligible, and the stability of HP is low. When [K^+^] is gradually increased to 150 mM (near physiological concentration), the states HP and I1 become stable (Fig. S4). However, the stability of states KC and I2 shows a modest increase. This indicates that even at high [K^+^], only secondary structures such as HP and I1 (have intra-chain contacts) are stable, and the tertiary structures such as I2 and KC (have inter-chain contacts) are unstable. Higher [K^+^] (*>* 150 mM) could stabilize the KC, but we restricted our investigation to [K^+^] = 150 mM, which is the physiological concentration.

To probe the effect of [Mg^2+^] on the dimerization pathway, we gradually increased [Mg^2+^] from 2 to 8 mM with the fixed background of [K^+^] = 150 mM (Fig. 3). We observed that even at low [Mg^2+^] (= 2 mM), the stability of KC increased significantly. From [Mg^2+^] = 4 to 8 mM, the KC becomes the most stable state. With increasing [Mg^2+^], the HP and I1 populations gradually decrease and shift towards the KC (Fig. 3). Interestingly, the population of the I2 state slightly increases up to [Mg^2+^] = 4 mM and decreases upon further increase in [Mg^2+^]. This indicates that [Mg^2+^] = 4 mM is the optimum ion concentration, where I2 is most stabilized. Since I2 has an unfolded chain, higher [Mg^2+^] (*>* 4 mM) shifts the equilibrium towards the structured KC state.

**Figure 3.**
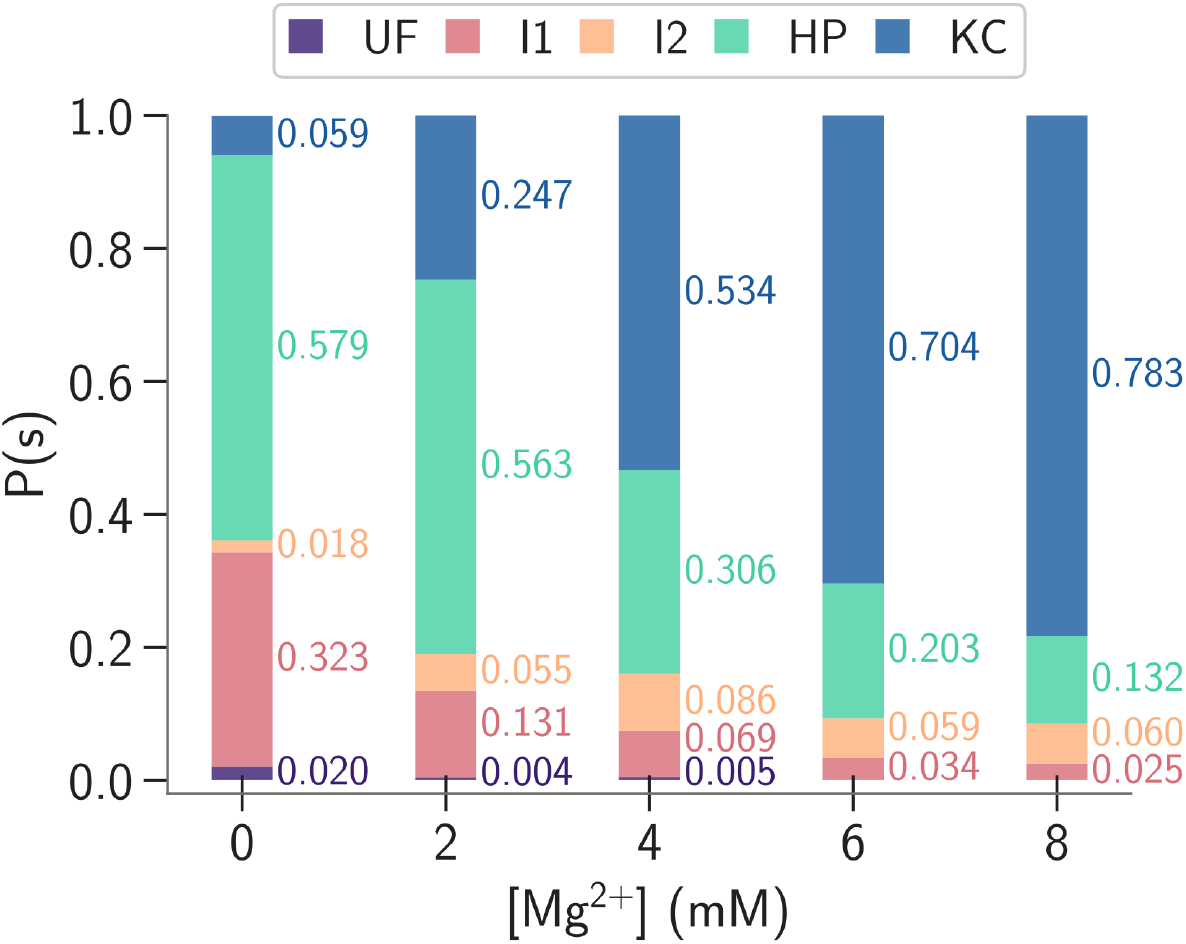
Probabilities of different states *s* (*s* = UF, I1, I2, HP, and KC), *P* (*s*), at different [Mg^2+^]. We excluded the ED state in the computation of equilibrium populations, as it is a sink. We utilized the K-means clustering algorithm, as implemented in Deeptime, ^38^ to calculate the probabilities.

All the trajectories collectively show a network of six states in the dimerization process (Fig. 2d). Initially, the network from HP bifurcates into two paths – one goes to KC and the other to I1. The KC transitions into the ED via the I2 intermediate (Fig. 2d). The network bifurcates again from the I1 state, where one path goes to the I2 state and the other to the UF state. The ED can form in two ways, either from the UF state or the I2 state (Fig. 2d). All the transitions are reversible except for the transitions from UF → ED and I2 → ED.

In the absence of Mg^2+^ and for [K^+^] = 50, 100, and 150 mM, all the trajectories reach ED only via UF (Fig. 2d, S5, Movie S1). The path preference changed in the presence of Mg^2+^. At [Mg^2+^] = 2 mM, ≈ 43% of the trajectories proceed to the ED state from UF, and ≈ 57% originate from I2 (Fig. 2d). At [Mg^2+^] = 4 mM, UF→ED transitions decreased to ≈ 29% and I2→ED transitions increased to ≈ 71%. From [Mg^2+^] = 6 mM and higher, all the trajectories proceed via I2→ED (Fig. 2d). Therefore, in this network, there can be several different paths for the dimerization process from HP to ED, depending on the [K^+^] and [Mg^2+^]. In the presence of only K^+^ (no Mg^2+^), the structures KC and I2 with inter-chain contacts are not stabilized. Hence, the preferred path for dimerization involves unfolded structures (I1 and UF), and is HP→I1→UF→ED (Fig. 2d). In contrast, with increasing [Mg^2+^], KC and I2 with inter-chain contacts are stabilized and the preferred path shifts to HP→KC→I2→ED (Fig. 2d, S6 and Movie S2). At low [Mg^2+^] (= 2 mM), we also observed two less frequent paths for ED formation – HP→I1→I2→ED and HP→KC→I2→I1→UF→ED (Fig. 2d). These two paths consist of both folded and unfolded states, indicating the critical point at which the dominant path switches. The force-pulling experiments on KC suggest that breaking the kissing-loop interactions is the slowest process, and increasing [K^+^] and [Mg^2+^] significantly strengthens the kissing-loop interactions.^16^ These experimental observations also substantiate our observation of the KC to ED transition via I2 without breaking the stable kissing-loop interactions at elevated [Mg^2+^].

## KC Populates an Ensemble of Bent Conformations and Transitions to ED via a T-shaped Semi-folded Intermediate

Previous smFRET studies revealed that the transition from the HP to the ED occurs via a bent-KC intermediate.^9^ To identify the bent conformations of the KC, we computed the bending angle (*θ*) (Fig. 4a) (see Methods). The distributions of *θ, P* (*θ*) as a function of [Mg^2+^] for all the KC conformations show a monomodal distributions with the mode ranging 138^∘^ ([Mg^2+^] = 0 mM) to 148^∘^ ([Mg^2+^] = 8 mM) (Fig. 4a). Although the peak position in the distributions shifts to higher *θ* with increasing [Mg^2+^], the most probable KC conformations remain bent. However, the question from the smFRET conclusions regarding how the hydrogen bond network rearranges from the bent-KC to the ED remains unclear.

**Figure 4.**
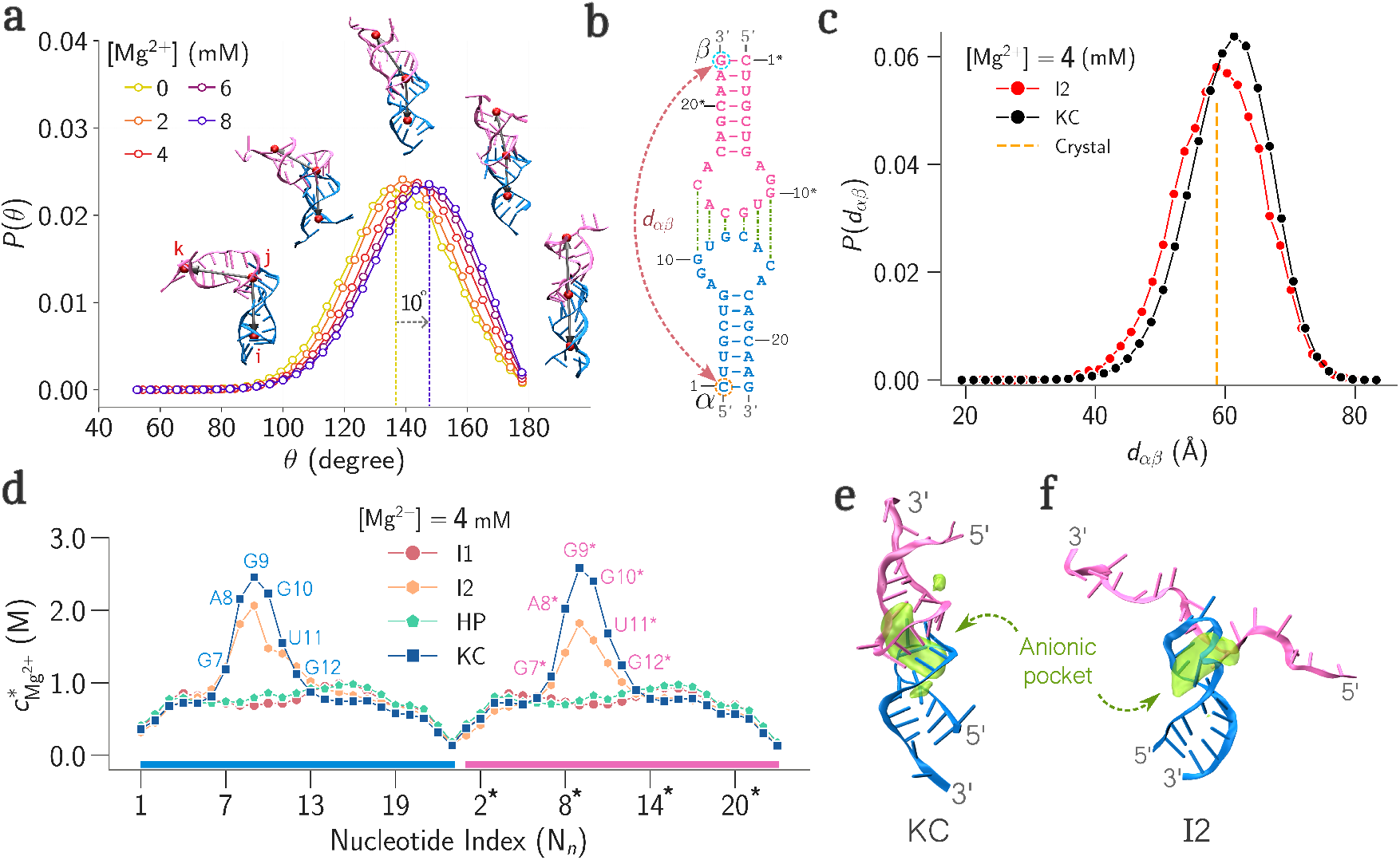
**a.** Probability distribution of the bending angle *θ, P* (*θ*), at different [Mg^2+^] (see Methods). The modes of *P* (*θ*) at [Mg^2+^] = 0 and 8 mM are highlighted with dotted lines. The representative structures with different *θ* are shown using the red beads (*i, j, k*) used for the *θ* calculation. **b**. The secondary structure of KC, highlighting the terminal residues (*α* and *β*) from DIS-1 and DIS-2 used for the smFRET studies.^9^ **c**. The distribution of the distance between *α* and *β* (*d*_*αβ*_) for states I2 and KC. The dashed line shows the corresponding *d*_*αβ*_ in the KC crystal structure (PDB ID: 1XPF^15^). **d**. The local Mg^2+^ concentration, 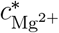for states I1, I2, HP, and KC. **e**. Spatial distribution of Mg^2+^ projected on a KC structure. **f**. Spatial distribution of Mg^2+^ projected on a I2 structure.

Our simulations in this work show that during the transition from KC to ED, one of the hairpin moieties of KC unfolds, keeping the kissing-loop hydrogen bonds intact to form the I2 state, and then the second hairpin unfolds and immediately forms the ED (Fig. 3d,4e,4f). In the experiments,^9^ the donor and the acceptor smFRET markers are labeled at the 5^′^ end of DIS1 (*α*) and the 3^′^ end of DIS2 (*β*), respectively (Fig. 4b). When the donor and acceptor FRET markers are at the termini, the KC and I2 states can give a similar FRET signal. To demonstrate this, we computed the distribution of the distances between *α* and *β* (*d*_*αβ*_) for the conformations in the KC and I2 states, and the distributions are similar (Fig. 4c). As a result, the I2 state is not identified in the experiments. The smaller 23nucleotide DIS construct undergoes dimerization without the need for any external proteins. In contrast, larger DIS constructs (≥ 35-nucleotides) require nucleocapsid proteins such as NCp7 to dimerize.^9,39–41^ For larger stem-loops, the population of I2-like intermediate would be unlikely due to the high energy barrier for unzipping under physiological [Mg^2+^] and temperature conditions. Hence, in the dimerization of larger stem-loops to form ED in HIV, the nucleocapsid proteins act as chaperones.^9,39–44^

## Mg^2+^ Condensation at the Anionic Pocket Stabilizes KC and I2 States

We computed the local Mg^2+^ concentration 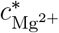 around the phosphate backbone of KC, HP, I2, and I1 states to probe the role of Mg^2+^ condensation in stabilizing these states (Fig. 4d) (see Methods). The 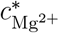 profile for I1 and HP does not show any significant peaks. In contrast, the KC and I2 states showed prominent 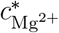 spikes, indicating a strong Mg^2+^ condensation in the vicinity of the nucleotides G7−G12 and G7^∗^−G12^∗^. The slight decrease in the 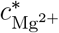 for I2 compared to KC is attributed to the presence of the unfolded hairpin in I2.

The nucleotides G7−G9 and G7^∗^−G9^∗^ are adjacent to the kissing loop region. Among these, nucleotides A8 and G9 are found to be bulged out in the crystal structures (Fig. 1b). These purines are dynamic in nature, showing bulge-in and bulge-out conformations, and are extensively studied using experiments ^10,15,22^ and atomistic simulations^11,25,26^ (discussed in the next section). The spike in 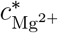 indicates that Mg^2+^ condensation can significantly influence the orientation of the bulged nucleotides (A8 and G9) (Fig. 4d). Further, the nucleotides G10−G12 and G10^∗^−G12^∗^ are directly involved in the formation of kissing-loop hydrogen bonds in both the KC and I2 states, which suggests that Mg^2+^ binding at the kissing loop region stabilizes the KC and I2 structures.

We computed the electrostatic potential surface of KC and I2 conformations using the PBEQ solver^45^ implemented in CHARMM-GUI^46^ (see Data Analysis in the SI). The potential surface shows that both conformations possess anionic cavities near the kissing-loop region, and the cavity in KC is larger than that of I2 (Fig. S7). Previous studies on KC also suggested the existence of the anionic pocket. ^25^ To visualize the specific Mg^2+^ binding on the RNA, we computed the average spatial density of Mg^2+^ ions around the KC and I2 structures (see Data Analysis in SI). The spatial density map confirms that the Mg^2+^ ions preferentially condense within the anionic pocket at the kissing-loop region (Fig. 4e,f). Extensive studies have investigated the role of divalent ions in stabilizing the RNA tertiary structures. ^32,34,47–54^ Previous studies suggest 1 mM Mg^2+^ is roughly comparable with 80 mM K^+^ in terms of charge screening capability. ^34,55–57^ Thus, K^+^ alone cannot stabilize the pocket, resulting in a lower population of KC at physiological levels of [K^+^] (≈ 150 mM). Since Mg^2+^ efficiently neutralizes the negatively charged cavity, a few mM Mg^2+^ within the physiological concentration range increases the KC population by stabilizing the anionic pocket.

The anionic pocket and the major groove-like geometry near the kissing-loop region have been exploited as a binding site for positively charged drug molecules. 4,5-disubstituted 2-deoxystreptamine (DOS) derivatives such as Neomycin, Paromomycin, Lividomycin, are positively charged molecules at pH = 7.4 and are reported to bind KC at this site.^35,58,59^ The binding affinities of these aminoglycoside derivatives are not solely dependent on non-specific electrostatic interaction. ^35^ They also form specific contacts with the sugar-phosphate backbone motif of KC for better target recognition.^35,60^ However, these drugs are not approved for HIV treatment as they have poor selectivity and bind to human RNAs and ribosomes. They are also toxic at concentrations where they show antiviral effects. Hence, there is a need to design derivatives of these drugs that exhibit high specificity and bind potently at low concentrations to the KC.

## Population of Kissing-complex (KC) Intermediate During Dimerization is Inherent to the DIS

The CG TIS RNA model^32,61^ we used is a native-centric model that requires information about the hydrogen bonds and tertiary stacking interactions present in the folded RNA structures. In this model, the parameters from an ideal A-form RNA are used for the standard canonical Watson-Crick pairs, and parameters extracted from the crystal structure are used for non-canonical hydrogen bonds. ^61^ To model the transition from HP to ED using the TIS RNA model, we need information about two sets of hydrogen bond networks − (i) the hydrogen bond network in the two HPs and (ii) the hydrogen bond network in the ED. All the hydrogen bonds used in the simulations are shown in Table S2. It is important to note that we did not use any hydrogen bond network information from any of the KC crystal structures.

The simulations in this work show that KC is a critical intermediate in the HP to ED transition (Fig. 2a-d). To quantify the similarity of KC structures populated in the simulations to the KC crystal structure, we computed the average contact map of the KC conformations obtained from the simulations and compared it with the contact map of the KC crystal structure (PDB ID: 1XPF) (Fig. S8). The average contact map of the KC conformations from simulations (Fig. S8, upper triangle) is in excellent agreement with the crystal contact map (Fig. S8, lower triangle). This observation suggests that the G10-C15 sequence in the DIS, which is involved in the KC formation, is also involved in canonical hydrogen bonding in ED with the second chain (Fig. 1b,c). In the case of an HP, the KC-forming sequence (G10-C15) is inside a 9-nucleotide-long flexible loop region (A8-A16) (Fig. 1b), and it interacts with the complementary sequence from another HP to form the KC complex.

## Dynamic Bulging of the Purine Bases Leads to Multiple Sub-ensembles of KC Conformations

Base-flipping is a vital mechanism in RNA-involved cellular processes, where specific nucleotides dynamically flip out of the central RNA helix, generating conformations that allow them to interact with proteins, enzymes, or other RNA molecules.^1,62–64^ The KC crystal structure showed two nucleobases in DIS-1 (A8 and G9) and DIS-2 (A8^∗^, and G9^∗^) hairpin motifs are bulged outwards and are known as bulged purines (Fig. 1b, 5a-b, S9a-b) (PDB ID: 1XPF^15^). However, multiple NMR studies reported different orientations, especially the bulged-in conformation, of these bulged purines. ^10,23,24^ Extensive atomistic simulations using different all-atom force fields have reported the dynamic nature of these bulged purines and suggested that the bulged-in conformations are dominant.^11,25,26^ The nucleotide G7 forms a strong Watson-Crick base pairing with C17, resulting in a fixed inward orientation of G7. Thus, we used G7 as a reference and classified the relative orientation of A8 and G9. Thus, to monitor the base bulging in the KC, we computed the pair distances between the base beads of (i) G7 and A8 (*d*_78_) and (ii) A8 and G9 (*d*_89_). We projected the 2D FES on the *d*_78_ and *d*_89_ along *x* and *y* axes, respectively, for DIS-1 (Fig. 5c and S10). The FES exhibits multiple minima, indicating distinct sub-ensembles of

**Figure 5.**
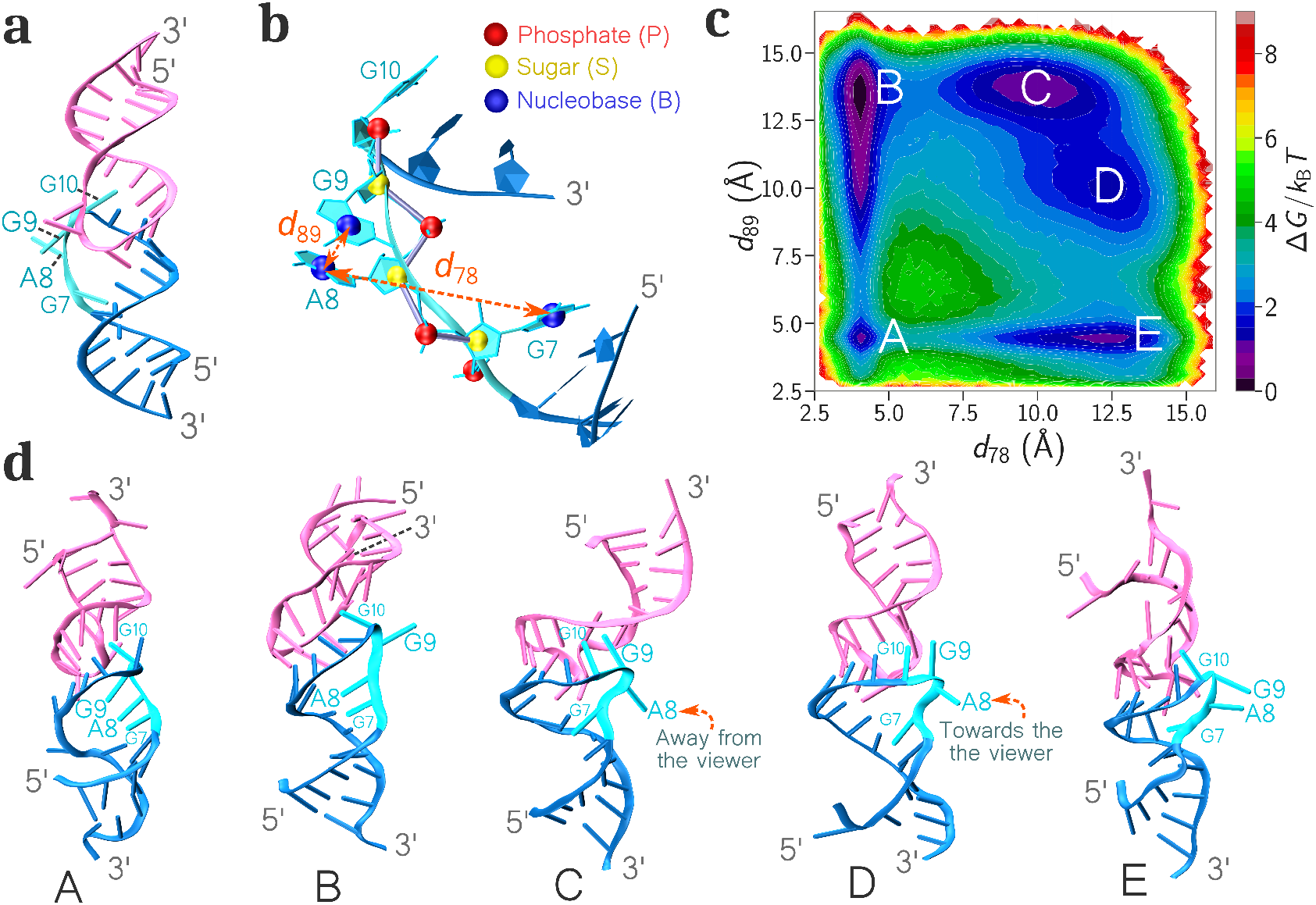
**a.** KC crystal structure (PDB ID: 1XPF). **b**. Distances between CG base beads (*d*_78_ and *d*_89_) used to monitor the relative orientation of the bulged nucleotides (A8 and G9) in DIS-1. **c**. 2D FES projected on *d*_78_ and *d*_89_ along *x* and *y* axes. **d**. Representative structures from basins A-E.

KC conformations with specific orientations of bulged purines (A8 and G9), which are labeled as states A to E. The representative structures are shown in Fig. 5d. Note that although the TIS-RNA model does excellent conformational sampling, it does not offer atomistic resolution of structures, as in all-atom simulations, due to its reduced degrees of freedom. Therefore, our claims of structural resemblance are qualitative. The conformations in state A (*d*_78_ ≈ 4.1 Å and *d*_89_ ≈ 4.5 Å) have both A8 and G9 bases bulged in (Fig. 5d). The structures from state A resemble the bulged-in structures reported in the NMR studies of the Lai variant of HIV-1 (which has A9 in Lai instead of G9, unlike the Mal variant) (PDB ID: 2F4X).^10,24^ The atomistic simulation studies have also shown that residues A8 and G9 mostly remain fully bulged in, which resembles the structures from state A.^11^ In state B (*d*_78_ ≈ 4.1 Å and *d*_89_ ≈ 13.5 Å) A8 is bulged in and G9 is bulged out (Fig. 5d). Havrila et. al also identified this broad ensemble in their atomistic simulations.^11^ States C (*d*_78_ ≈ 10.0 Å and *d*_89_ ≈ 13.7 Å) and D (*d*_78_ ≈ 12.5 Å and *d*_89_ ≈ 10.0 Å) are broad adjacent basins, where A8 is bulged out and G9 points towards the kissing-loops, forming a consecutive stack with G10. However, states C and D differ in the orientation of A8 base as shown in Fig. 5d. These conformations partially resemble the structure suggested by the NMR studies by Mujeeb et al. on the Lai variant (PDB ID: 1BAU ^22^), where A8 remains bulged in and A9 forms a consecutive stack with G10, pointing towards the kissing-loop. However, A8 is bulged out in states C and D in our simulations. In state E (*d*_78_ ≈ 12.5 Å and *d*_89_ ≈ 4.5 Å), both A8 and G9 are bulged out. The structures with bulged-out G9 and A8 were reported in the crystal structures of KC. ^15,21^ We observed similar results for bulged purines in DIS-2 (A8^∗^ and G9^∗^) (Fig. S9a-g). Thus, our simulations sampled most of the conformational states reported by different experimental techniques for the bulged purines of the KC, and we further show that they exist in dynamic equilibrium. We should mention that although we qualitatively sample various sub-ensembles of KC conformations involving the bulged purines, we cannot quantitatively capture their relative populations as they depend on the dihedral potentials, and in the TIS RNA model, they are parameterized for A-form RNA helix.

The mechanism of HIV-1 RNA dimerization pathways is essential for understanding the genome packaging process associated with the retroviral assembly. In this study, we present a comprehensive assembly mechanism for the dimerization of two DIS hairpins, involving multiple intermediates that form an equilibrium network before irreversibly forming the ED. Among all the transitions between the intermediate states, the transition from KC to ED via I2 shows the key rearrangement of the hydrogen bond network, and this completes the mechanism proposed from the previous smFRET experiment.^9^ The dominant pathway in the network leading to ED formation in the presence of divalent ions such as Mg^2+^ is through the KC. Mg^2+^ ions condense in the anionic cavity of the KC and stabilize this intermediate. The stability of the KC decreases in the sole presence of monovalent ions such as K^+^ present at physiological levels, thereby shifting the dominant dimerization pathway to one that is devoid of KC. The dynamics of bulged purines in the KC lead to the population of different sub-ensembles of KC conformations, reconciling the KC structures from the previous crystallography, NMR, and simulation studies. ^10,11,15,21,22,24–26^ Previous studies have suggested that these dynamic purines may play a critical role in NCp7 binding, correlating this with the activity of NCp7 in DIS dimerization.^41,44^ Additionally, the anionic pockets in both KC and I2 make these structures potential drug targets. Although aminoglycoside derivatives showed significant potential to halt the dimerization process at KC, due to toxicity and reduced inhibition activity, this potential has not yet been fully realized.^1,35,60^

This study complements the existing literature and provides insights into the implications for complex RNA-RNA interactions involved in bioassemblies. Although this study presents the overall free-energy landscape of the dimerization process, the use of a small 23-nucleotide DIS construct as the model system limits its scope. Further studies should be on longer stemloops and involve nucleocapsid proteins that are known to participate in the dimerization process^9,42–44,65^ for a broader perspective.

## Methods

### Coarse-Grained MD Simulations

We used the three interaction site (TIS) RNA model^32,61^ to investigate the dimerization of RNA HPs to form the ED. In the TIS-RNA model, a nucleotide is represented by three beads positioned at the center of mass of the phosphate group (P), the sugar (S), and the nucleobase (B). All CG simulations were carried out using the OpenMM molecular simulation toolkit.^66^ TIS model details and the simulation protocols are in the SI.

### Free Energy Surface (FES) from CG Simulations

The thermodynamic states populated during the dimerization process were identified by computing the 2D free energy surface (FES), *G*(*C*_1_, *C*_2_), projected onto two collective variables (*C*_1_ and *C*_2_). We computed the *G*(*C*_1_, *C*_2_) using the following equation,

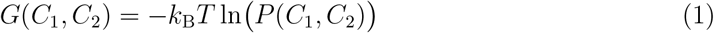

where *P* (*C*_1_, *C*_2_) is the joint probability distribution of *C*_1_ and *C*_2_ at a given [Mg^2+^] and *k*_B_ is the Boltzmann constant. For the classification of the overall states, we used 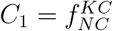 and 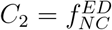 (Fig. 2a-c). For the classification of KC conformational sub-ensembles populated due to the bulging of purine bases, we used *C*_1_ = *d*_78_ and *C*_2_ = *d*_89_ for DIS-1 (Fig. 5b-c) and 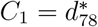 and 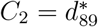for DIS-2 (Fig. S9b-g).

### Bending Angle Calculation

To calculate the bending angle (*θ*) of a KC conformation, we used three center of mass (COM) coordinates *i, j*, and *k* shown as red beads in representative KC structures (Fig. 5). For calculating the COM, we used backbone sugar beads of nucleotides U4, U5, A19, and A20 for *i*; nucleotides U11, G12, C13, U11^∗^, G12^∗^, and C13^∗^ for *j*; nucleotides U4^∗^, U5^∗^, A19^∗^, and A20^∗^ for *k*.

## Supporting Information (SI)

Simulation details and additional data analysis; Parameters and the hydrogen bond network used in the CG simualtions (Table S1 and S2); Representative trajectories at different [Mg^2+^] (Fig. S1, S5, and S6); The 2D FES projected on 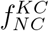 and 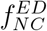for Mg^2+^ and K^+^ concentrations (Fig. S2 and S3); Equilibium probabilities for states at different [K^+^] (Fig. S4); The electrostatic potential surface for KC and I2 (Fig. S7); The average contact map of KC from simulation with contact map from the crystal structure (PDB ID; 1XPF) (Fig. S8); The 2D FES projected on base-base distances of bulged purines at different [Mg^2+^] for DIS-2 (Fig. S9) and in DIS-1 (Fig. S10).

## Supporting information

Methods, Data Analysis and Supporting Tables and Figures

## Acknowledgement

GR acknowledges funding from the Science and Engineering Research Board (SERB) through the grant CRG/2023/002817. DM acknowledges the research fellowship from the Indian Institute of Science, Bangalore. SH acknowledges the Prime Minister Research Fellowship (PMRF). We acknowledge the National Supercomputing Mission (NSM) for providing computing resources of “Param Pravega” at IISc and “Param Brahma” at IISER Pune, supported by the Ministry of Electronics and Information Technology (MeitY) and the Department of Science and Technology (DST), Government of India.

## Notes

### Competing Interest Statement

The authors have declared no competing interest.

