## Supplementary material for "Dimerization Mechanism of HIV-1 RNA Hairpins to Extended Duplex Structures": Methods, Data Analysis and Supporting Tables and Figures

### Supporting Information for "Dimerization Mechanism of HIV-1 RNA Hairpins to Extended Duplex Structures"

### Coarse-grained (CG) simulations

#### Three-Interaction-Site (TIS) Model

We performed the CG simulations using the three interaction site (TIS) RNA model.[1,2] In the TIS model, a nucleotide is represented by three beads positioned at the center of mass of the phosphate group (P), the sugar (S), and the nucleobase (B). The TIS energy function includes potentials to model bond length ($U_{\mathrm{BL}}$), bond angle ($U_{\mathrm{BA}}$), excluded volume interactions ($U_{\mathrm{EV}}$), stacking between consecutive bases ($U_{\mathrm{ST}}$), tertiary stacking (stacking between non-consecutive bases) ($U_{\mathrm{TST}}$), electrostatic interactions ($U_{\mathrm{EL}}$) and native hydrogen bond ($U_{\mathrm{HB}}$) interactions present in the folded structure. The energy function[1] is given by

$$U_{\mathrm{TIS}} = U_{\mathrm{BL}} + U_{\mathrm{BA}} + U_{\mathrm{EV}} + U_{\mathrm{EL}} + U_{\mathrm{ST}} + U_{\mathrm{HB}} + U_{\mathrm{TST}} \tag{S1}$$

The bond length and bond angle interactions are modeled using harmonic potentials given by,

$$U_{\mathrm{BL}} = k_\rho(\rho - \rho_0)^2 \tag{S2}$$

$$U_{\mathrm{BA}} = k_\alpha(\alpha - \alpha_0)^2 \tag{S3}$$

where $\rho_0$ and $\alpha_0$ are the bond distance and bond angles at equilibrium from the ideal A-form RNA helix. The $k_\rho$ values are 23, 64, and 10 in kcal.mol$^{-1}$·Å$^{-2}$ for P→S, S→P (' →' implies 5′→3′) and S−B bonds, respectively. The $k_\alpha$ values are 5 kcal mol$^{-1}$ rad$^{-2}$ for angles involving the base beads, and 20 kcal mol$^{-1}$ rad$^{-2}$ otherwise.

The potential used for modeling the RNA hydrogen bonding network is given by,

$$U_{\mathrm{HB}} = U_{\mathrm{HB}}^0 \exp(-u) \tag{S4}$$

where,

$$u = 5.0(r-r_0)^2+1.5\{(\theta_1-\theta_{1,0})^2+(\theta_2-\theta_{2,0})^2\}+0.15\{(\psi_0-\psi_{0,0})^2+(\psi_1-\psi_{1,0})^2+(\psi_2-\psi_{2,0})^2\}.$$

The value of $U^0_{\rm HB}$ is -2.9 kcal·mol$^{-1}$. The parameters $r_0$, $\theta_{1,0}$, $\theta_{2,0}$, $\psi_{0,0}$, $\psi_{1,0}$, and $\psi_{2,0}$ for the canonical hydrogen bonds are calculated from the coarse-grained model of the ideal A-form RNA helix and for the noncanonical hydrogen bonds are calculated from the coarse-grained model of the crystal structure (PDB ID: 462D).[1–3] The canonical and noncanonical hydrogen bonds used in the force field are shown in the secondary structures (see Fig. 1b,c in the Main text). The definitions of $r$, $\theta_1$, $\theta_2$, $\psi_0$, $\psi_1$, and $\psi_2$ are described in the work of Denesyuk et. al.[2]

The stacking interactions between two consecutive nucleotides are sequence-dependent ($U_{\rm ST}$) and are given by

$$U_{\rm ST} = \frac{U^0_{\rm ST}}{1 + k_r(r-r_0)^2 + k_\phi(\phi_1-\phi_{1,0})^2 + k_r(\phi_2-\phi_{2,0})^2} \tag{S5}$$

where, $k_r$ is 1.4 Å$^{-2}$ and $k_\phi$ is 4.0 rad$^{-2}$. The $r_0$, $\phi_{1,0}$, and $\phi_{2,0}$ are calculated from the coarse-grained structure of an ideal A-form RNA helix.[1,2,4] The values of $U^0_{\rm ST}$ and all the definitions of $r$, $\phi_1$, and $\phi_2$ are defined in the original work of Denesyuk et al.[1]

The tertiary stacking interaction between two non-adjacent nucleotide bases ($U_{\rm TST}$) is given by

$$U_{\rm TST} = \frac{U^0_{\rm TST}}{1+u} \tag{S6}$$

where,

$$u = 5.0(r-r_0)^2+1.5\{(\theta_1-\theta_{1,0})^2+(\theta_2-\theta_{2,0})^2\}+0.15\{(\psi_0-\psi_{0,0})^2+(\psi_1-\psi_{1,0})^2+(\psi_2-\psi_{2,0})^2\}.$$

The value of $U^0_{\rm TST}$ is taken to be -6.5 kcal·mol$^{-1}$. The definitions of the other variables in the potential are the same as the hydrogen bonding interaction potential used in the work of Denesyuk et al.[2]

The electrostatic interaction between the phosphate beads is computed using the Coulomb potential given by.

$$U_{\mathrm{EL}} = \frac{Z_i Z_j}{4\pi\epsilon_0\epsilon_r r_{ij}} \tag{S7}$$

The dielectric constant of water is a function of temperature ($\epsilon_r(T)$) and is given by

$$\epsilon_r(T) = 87.740 - 0.4008T + 9.398 \times 10^{-4}T^2 + 1.410 \times 10^{-6}T^3 \tag{S8}$$

A modified LJ potential is used to model the excluded volume interactions between the beads and is given by

$$U_{\mathrm{EV}} = \begin{cases} \epsilon_{ij}\left[\left(\frac{d_s}{r + d_s - d_{ij}}\right)^{12} - 2\left(\frac{d_s}{r + d_s - d_{ij}}\right)^{6} + 1\right] & \text{if } r \leq d_{ij} \\ 0 & \text{if } r > 0 \end{cases} \tag{S9}$$

where $d_s$ is the diameter of the smallest ion present in the system, $d_{ij} = R_i + R_j$, $\epsilon_{ij} = \sqrt{\epsilon_i\epsilon_j}$. The values of $R_i$ and $\epsilon_i$ for phosphate (P), sugar (S), and bases (A, G, C, and U) are taken from the work of Denesyuk et al.[2] The values of $R_i$ and $\epsilon_i$ for the ions are given in Table S1.

#### CG Simulation Details

We used the TIS-RNA[1,2] model to study the dimerization of the DIS hairpin's (HPs) to extended duplex (ED). The initial conformations for the simulations were generated using the ED crystal structure.[3] We converted the ED structure into the CG representation and unfolded it by performing a simulation at $T$ = 350 K and [$K^+$] = 150 mM. To expedite the unfolding, we switched off the hydrogen bond potential between the bases in this simulation.

To generate the initial HP conformations for the simulation, we performed a simulation with two CG unfolded chains at $T$ = 300 K, [$Mg^{2+}$] = 4 mM and [$K^+$] = 150 mM. In this simulation, the hydrogen bond potential between the bases that facilitate HP formation was switched on, while the hydrogen bond potential between the bases that facilitate ED formation was switched off to expedite the HP formation. This protocol generates initial conformation with two HPs randomly oriented relative to each other. We used the conformations where the two HPs are at least 15 Å apart from each other as initial conformations for production runs. For all the simulations (except those for initial structure generation), the hydrogen bond potential between the bases that support HP and ED formation was simultaneously active.

We performed Langevin dynamics simulations using OpenMM[5] at $T$ = 333 K using a cubic box of length 150 Å. The water solvent is implicitly modeled. Simulations were performed at various ion concentrations. Initially, the simulations were performed in $[K^+]$ = 50, 100, and 150 mM. Simulations were also performed at $[Mg^{2+}]$ = 2, 4, 6, and 8 mM, with a fixed background of $[K^+]$ = 150 mM. The $K^+$, $Mg^{2+}$, and $Cl^-$ ions were randomly inserted into the box to maintain the required concentration. We performed 7 independent simulation runs for each ion concentration. The simulation lengths are not the same for all the trajectories, as we stopped the simulations within a few $\mu$s after the formation of ED, which is irreversible. The durations of the trajectories varied from ≈6 $\mu$s to 20 $\mu$s. We discarded the initial 500 ns of the data from each CG trajectory for analysis.

The equation of motion for the particles in the simulation is given by

$$m_i \ddot{\vec{r}_i} = -m_i \gamma \dot{\vec{r}_i} + \vec{F}_i + \vec{\Gamma}_i \tag{S10}$$

where, $\vec{r_i}$ and $\gamma$ are coordinates and friction coefficient of the $i$th particle, respectively. The deterministic force on the particle $i$ is given by $\vec{F_i} = -\dfrac{\partial U_{\mathrm{TIS}}(\vec{r_i})}{\partial \vec{r_i}}$, and $\vec{\Gamma}_i$ is an uncorrelated random force with a white noise spectrum. The autocorrelation function of the random force in discretized form is $\left\langle \vec{\Gamma}(t) \cdot \vec{\Gamma}(t+nh) \right\rangle = \frac{2\gamma m_i k_{\mathrm{B}} T}{h} \delta_{0,n}$, where $n$ = 0, 1, ... and $\delta_{0,n}$ is the Kronecker delta function. We have used the *LangevinIntegrator* module in OpenMM to integrate the equations of motion. For better conformational sampling, we used a low friction coefficient ($\gamma = 1$ ps$^{-1}$) with an integration time step of 2 fs.

We applied a harmonic potential with a low force constant ($k_s = 2$ kcal mol$^{-1}$ Å$^{-2}$) between the center of mass of each DIS chain. This potential is activated only when the distance between the COMs exceeds the cutoff distance ($r_s^{\text{cut}} = 50$ Å) and remains inactive otherwise. $r_s^{\text{cut}}$ is approximately 2.5 times the radius of gyration of the ED ($R_g^{\text{ED}} \approx 19.3$ Å) or KC ($R_g^{\text{KC}} \approx 20.0$ Å) crystal structures, which ensures that there is enough distance between the two DIS chains in the unbound states (such as HP, UF and I1) and this potential does not bias the binding. The Particle Mesh Ewald (PME) algorithm was used to compute the long-range Coulomb interactions. We used the TIS2AA program[6] to generate all-atom representations of the coarse-grained structures and VMD[7] for visualization and representation of the conformations.

### Data Analysis

##### Fraction of native Contacts

We computed the fraction of native contacts (NC) between a set of nucleotides ($\lambda$) in a conformation $i$ using

$$f_{\text{NC}}^{\lambda,(i)} = \frac{N_{\text{NC}}^{\lambda,(i)}}{N_{\text{NC}}^{\lambda,(\text{cry})}} \tag{S11}$$

where $N_{\text{NC}}^{\lambda,\,\text{cry}}$ and $N_{\text{NC}}^{\lambda,\,(i)}$ are the number of native contacts present between the nucleotides belonging to set $\lambda$ in the crystal structure and the $i^{\text{th}}$ conformation, respectively.[8,9] If the distance between a pair of beads is less than the cutoff distance $r_{cut}$ (= 15 Å),[2,10] then that pair is considered to be in contact.

##### Computation of Contact maps

To compute the average contact map, the pair distances between all the RNA CG beads were calculated, excluding the beads that are not at least three nucleotides apart from each other in the RNA sequence. It is considered a contact between the given pair of beads if a pair distance is less than the cutoff distance $r_{cut}$ (= 15 Å).[2,10] We averaged over all the conformations in the ensemble of KC.

#### Local Ion Concentration Around RNA

We computed the local ion concentration ($c_j^*$) to investigate the condensation of positively charged divalent metal ions around phosphate beads using the equation[2]

$$c_j^* = \frac{1}{N_\mathrm{A} V_c} \int_0^{r_c} \mathrm{d}r \; 4\pi r^2 \; \rho_j(r) \tag{S12}$$

where, $\rho_j(r)$ is the number density of a specific type of ion $j$ at a distance $r$ from the specific RNA bead, $V_c$ is the spherical volume with a radius $r_c$, and $N_\mathrm{A}$ is the Avogadro number. The cutoff radius is defined as $r_c^{(\mathrm{CG})} = R_{\mathrm{M}^{n+}} + R_\mathrm{P} + \Delta r$ ($\approx 5$ Å), where $R_{\mathrm{M}^{n+}}$ is the radius of the the metal ion $\mathrm{M}^{n+}$ (Table S1), $R_\mathrm{P}$ (= 2.1 Å) is the CG radius of a phosphate bead[2] and $\Delta r$ (= 1.6 Å) is the margin distance.

#### Spatial Distribution of Ions Around RNA

We employed the volmap plugin in VMD software[11] to create spatial density maps depicting the distribution of $Mg^{2+}$ ions around RNA nucleotides. To begin, RNA beads from all conformations of a particular state were superimposed on a reference structure of the same state, and the positions of metal ions in each simulation frame were adjusted accordingly. The superimposition was performed using the Kabsch method,[12] as implemented in VMD. In our analysis, metal ions were represented as spheres with their respective van der Waals radii. A grid of 10 Å$^3$ was constructed to divide the space around the RNA nucleotides. Each grid point was marked as occupied (value of 1) if overlapped by a sphere representing a metal ion; otherwise, it was assigned a value of 0. These values were averaged across all frames of the trajectory. VMD was also used for structural superposition[13] and image rendering.[14]

Table S1: RNA[2] and ions[15,16] parameters used in the CG simulations.

| Type | $R_i$ (Å) | $m_i$ (amu) | $\epsilon_i$ (kcal mol$^{-1}$) | $Z_i$ |
|---|---|---|---|---|
| P | 2.1 | 62.971 | 0.2 | -1 |
| S | 2.9 | 131.108 | 0.2 | 0 |
| A | 2.8 | 134.119 | 0.2 | 0 |
| G | 3.0 | 150.118 | 0.2 | 0 |
| C | 2.7 | 110.094 | 0.2 | 0 |
| U | 2.7 | 111.079 | 0.2 | 0 |
| $Mg^{2+}$ | 1.353 | 24.305 | 0.009 | +2 |
| $K^+$ | 1.590 | 39.098 | 0.279 | +1 |
| $Cl^-$ | 2.760 | 35.453 | 0.012 | -1 |

Table S2: The hydrogen bonds used in the CG simulations. [a]Non-canonical hydrogen bonds.

| Hairpin (counts) | Extended duplex (counts) |
|---|---|
| **DIS-1:** C1–G23(3), U2–A22(2), U3–A21(2), G4–C20(3), C5–G19(3), U6–A18(2), G7–C17(3) **DIS-2:** C1*–G23*(3), U2*–A22*(2), U3*–A21*(2), G4*–C20*(3), C5*–G19*(3), U6*–A18*(2), G7*–C17*(3) | C1–G23*(3), U2–A22*(2), U3–A21*(2), G4–C20*(3), C5–G19*(3), U6–A18*(2), G7–C17*(3), G9–A16*(2)[a], G10–C15*(3), U11–A14*(2), G12–C13*(3), C13–G12*(3), A14–U11*(2), C15–G10*(3), A16–G9*(2)[a], C17–G7*(3), A18–U6*(2), G19–C5*(3), C20–G4*(3), A21–U3*(2), A22–U2*(2), G23–C1*(3) |

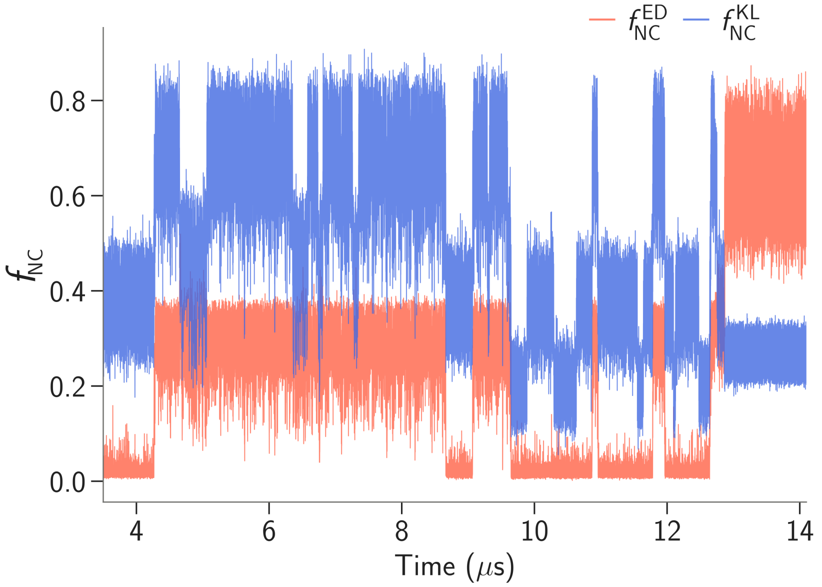


Figure S1: A representative trajectory showing the fraction of native contacts of ED ($f_{NC}^{ED}$) and the fraction of native contacts of KC ($f_{NC}^{KC}$) at $[Mg^{2+}] = 4$ mM and temperature 333 K.

‘

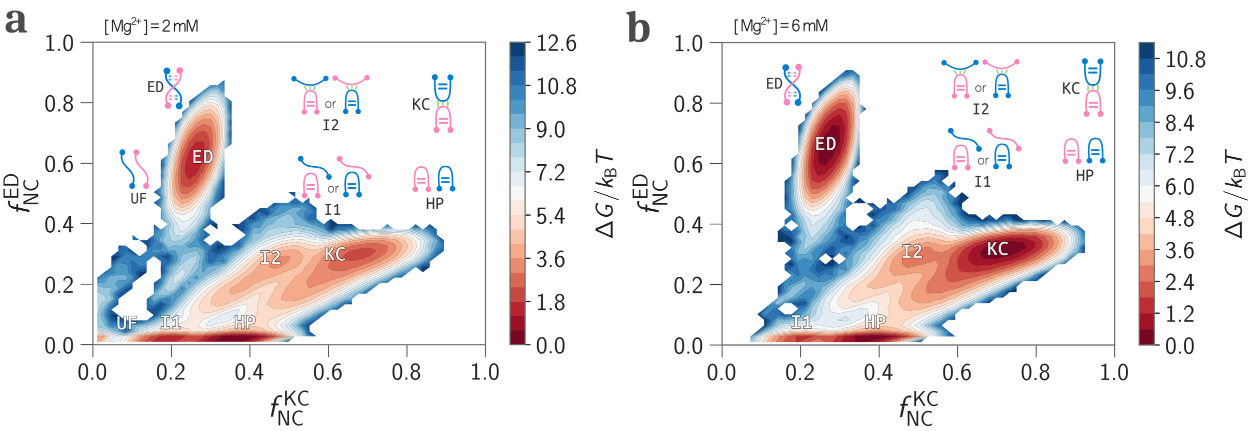


Figure S2: The 2D FES projected onto $f_{NC}^{KC}$ and $f_{NC}^{ED}$ for **a.** $[Mg^{2+}] = 2$ mM and **b.** $[Mg^{2+}] = 6$ mM. All have a fixed background $[K^+] = 150$ mM. The state ED is a sink. The schematic structures are shown to highlight the structural features in each state.

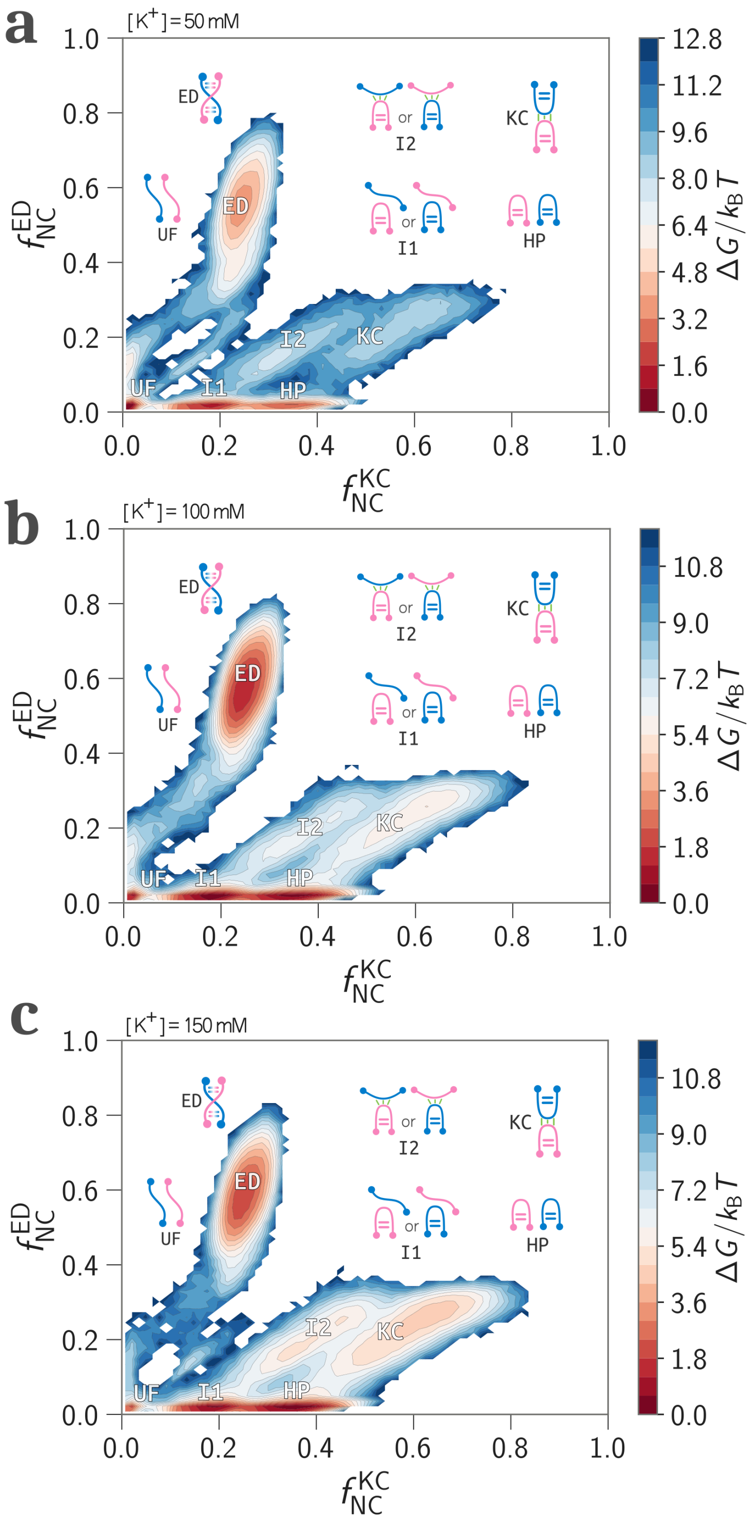


Figure S3: The 2D FES projected onto $f_{\rm NC}^{\rm KC}$ and $f_{\rm NC}^{\rm ED}$ for **a.** $[K^+]$ = 50 mM, **b.** $[K^+]$ = 100 mM, and **c.** $[K^+]$ = 150 mM. The state ED is a sink. The schematic structures are shown to highlight the structural features in each state.

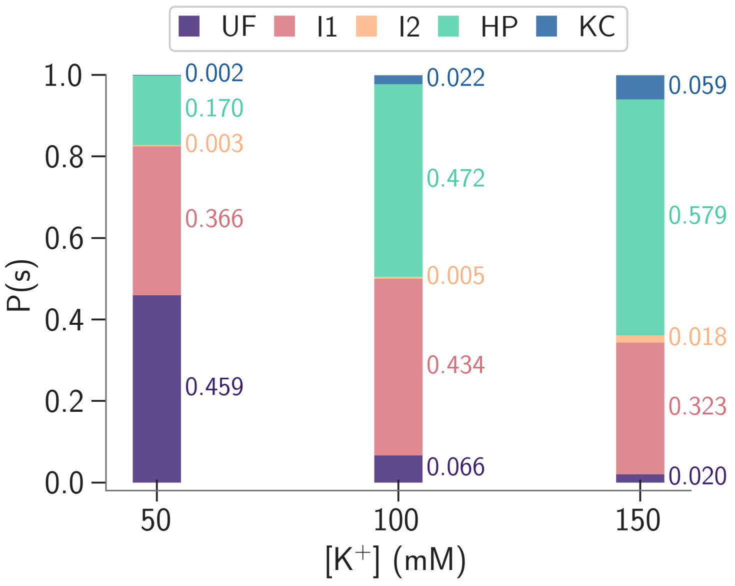


Figure S4: The equilibrium probabilities of different states $s$ (= UF, I1, I2, HP, and KC except ED), $P(s)$, at different [K$^+$].

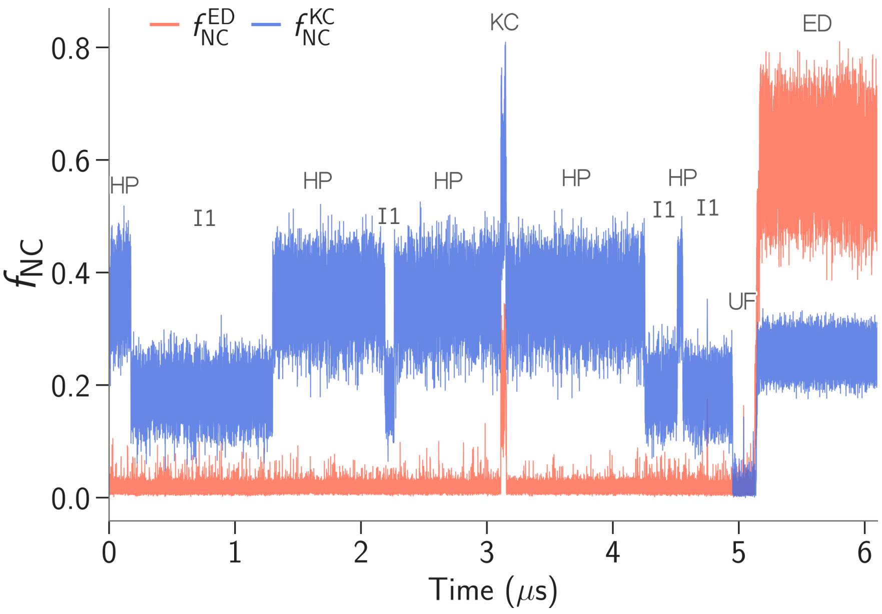


Figure S5: A representative trajectory showing transitions between different intermediate states before irreversibly forming the ED at [Mg$^{2+}$] = 0 mM, [K$^+$] = 150 mM and temperature 333 K.

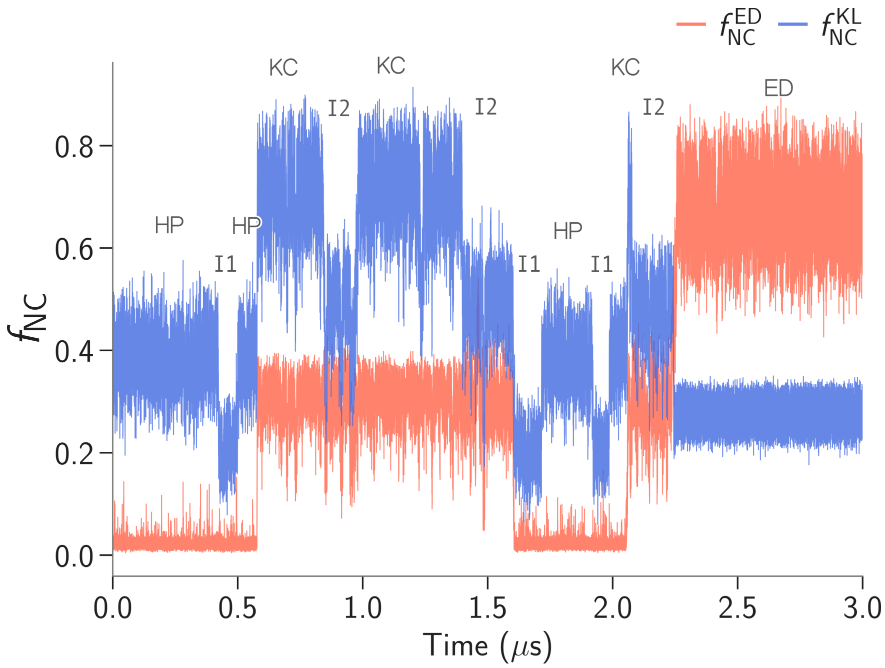


Figure S6: A representative trajectory showing transitions between different intermediate states before irreversibly forming the ED at [$Mg^{2+}$] = 8 mM, [$K^+$] = 150 mM, and temperature 333 K.

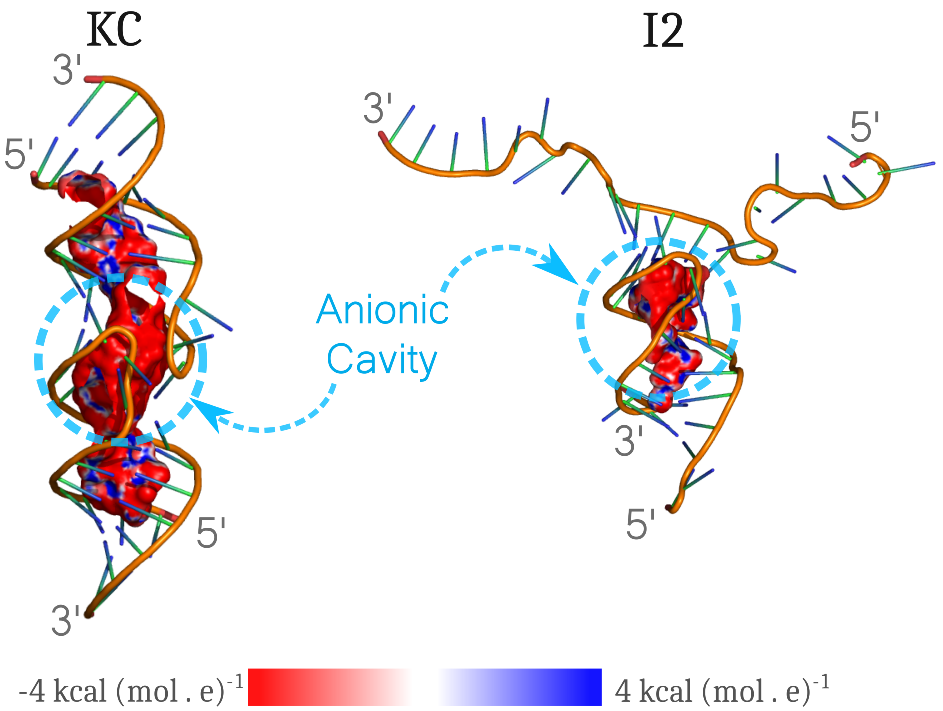


Figure S7: The anionic cavity at the kissing-loop region is highlighted using the dashed sky blue circle in KC (left) and I2 (right). The anionic cavity size is large in KC compared to I2. The electrostatic potential surfaces are calculated using the PBEQ Solver server[17] present in CHARMM-GUI[18] by solving the Poisson-Boltzmann equation.[19] The parameters used for the calculation: (i) dielectric constant for the reference environment, $\varepsilon_R = 1.0$, (ii) dielectric constant for the RNA interior, $\varepsilon_P = 1.0$, (iii) solvent dielectric constant, $\varepsilon_W = 80$, and (iv) salt concentration, [salt] $=$ 0.15 M. The electrostatic potential surface is projected on the cavity and visualized using the surface cavity module in PYMOL with cavity detection radius $= 7$ Å and cavity detection cutoff $= 4$ solvent radius.[20]

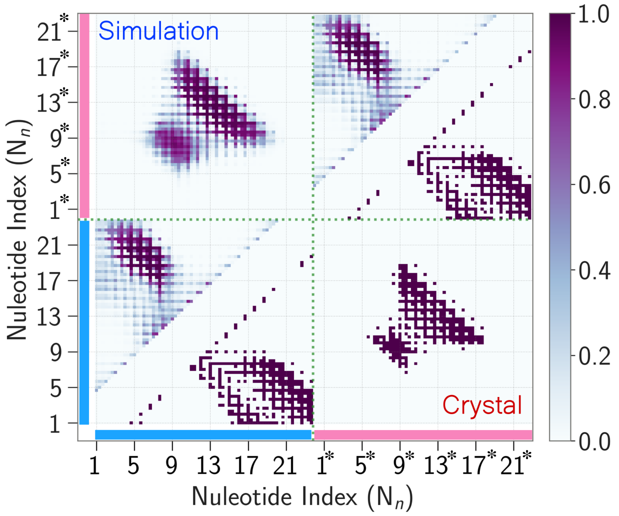


Figure S8: Comparison of the average native contact map of KC conformations obtained from the simulations (upper triangle) to the KC crystal structure (lower triangle)(PDB ID: 1XPF[21]).

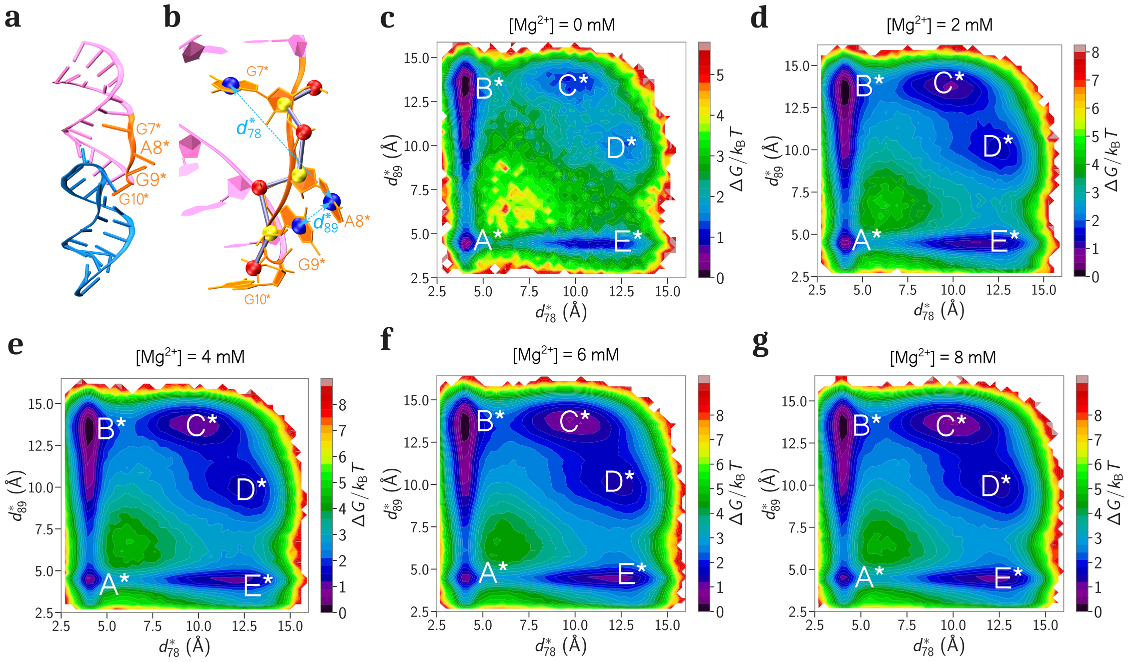


Figure S9: **a.** KC crystal structure (PDB ID: 1XPF). **b.** Distances between CG base beads ($d^*_{78}$ and $d^*_{89}$) is used to monitor the relative orientation of the bulged nucleotides (A8* and G9*) in DIS-2. 2D FES of the bulged purines (A8*, G9*) of DIS-2 projected on $d^*_{78}$ and $d^*_{89}$ along the $x$ and $y$ axes for **c.** $[Mg^{2+}] = 0$ mM, **d.** $[Mg^{2+}] = 2$ mM, **e.** $[Mg^{2+}] = 4$ mM, **f.** $[Mg^{2+}] = 6$ mM, and **g.** $[Mg^{2+}] = 8$ mM, with a fixed background of $[K^+] = 150$ mM.

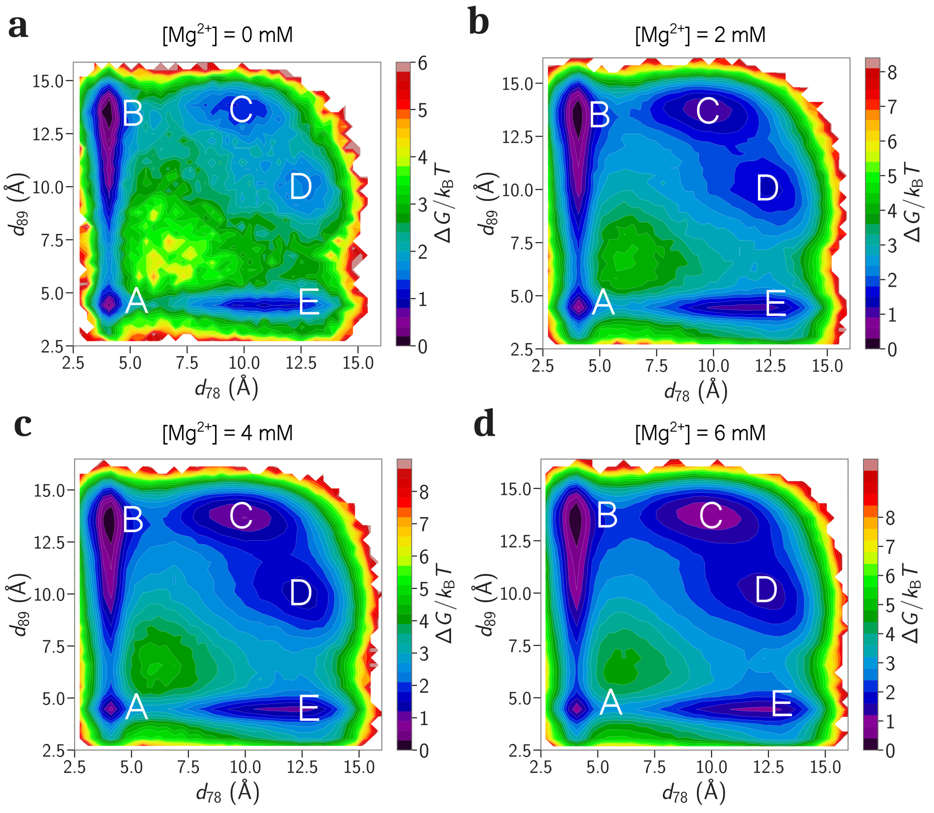


Figure S10: 2D FES's of the bulged purines (A8, G9) of DIS-1 projected on $d_{78}$ and $d_{89}$ along the $x$ and $y$ axes for **c.** $[Mg^{2+}] = 0$ mM, **d.** $[Mg^{2+}] = 2$ mM, **e.** $[Mg^{2+}] = 4$ mM, and **f.** $[Mg^{2+}] = 6$ mM, with a fixed background of $[K^+] = 150$ mM.

### Movie S1

This movie shows a transition from HP→I1→UF→ED.

### Movie S2

This movie shows a transition from HP→KC→I2→ED.
